# Proximal Molecular Probe Transfer (PROMPT), a new approach for identifying sites of protein/nucleic acid interaction in cells by correlated light and electron microscopy

**DOI:** 10.1101/2023.05.30.542936

**Authors:** Guillaume A Castillon, Sebastien Phan, Junru Hu, Daniela Boassa, Stephen R Adams, Mark H Ellisman

## Abstract

The binding and interaction of proteins with nucleic acids such as DNA and RNA constitutes a fundamental biochemical and biophysical process in all living organisms. Identifying and visualizing such temporal interactions in cells is key to understanding their function. To image sites of these events in cells across scales, we developed a method, named PROMPT for PROximal Molecular Probe Transfer, which is applicable to both light and correlative electron microscopy. This method relies on the transfer of a bound photosensitizer from a protein known to associate with specific nucleic acid sequence, allowing the marking of the binding site on DNA or RNA in fixed cells. The method produces a fluorescent mark at the site of their interaction, that can be made electron dense and reimaged at high resolution in the electron microscope. As proof of principle, we labeled *in situ* the interaction sites between the histone H2B and nuclear DNA. As an example of application for specific RNA localizations we labeled different nuclear and nucleolar fractions of the protein Fibrillarin to mark and locate where it associates with RNAs, also using electron tomography. While the current PROMPT method is designed for microscopy, with minimal variations, it can be potentially expanded to analytical techniques.

## INTRODUCTION

There are limited methods for visualizing the specific interaction of proteins-of-interest with DNA and RNA in cells by correlated light and electron microscopy (CLEM), that enable their identification, localization, and quantification of the bound fraction of protein to DNA/RNA. This is crucial for understanding the behavior of transcription factors, DNA or RNA polymerases, DNA repair enzymes, or mRNA editing enzymes. Potential applications include an ultrastructural snapshot of the interacting RNA binding proteins and mRNA in neuronal axons and dendritic spines that enable the spatially limited protein synthesis necessary for synaptic changes during neuronal plasticity. A key outcome and one goal of the project reported is to propel capabilities for localization of specific nucleic acid sequences without the inherent degradation of structure associated with methods now used for in situ hybridization – thus providing localizations of specific sequences in the context of the cell, but now with extremely well-preserved fine structure.

A plethora of methods is available for detecting and imaging protein-protein interactions using FRET between proteins tagged with fluorescent proteins(*1, 2*). Comparable methods for imaging protein-DNA/RNA interactions require fluorescent labeling of specific nucleotide sequences involved in their binding but the harsh conditions required for Fluorescent In Situ Hybridization (FISH) are incompatible with optimal cellular ultrastructure preservation for EM(*3*). FP-tagging for proteins and for RNA or DNA can be challenging as overexpression masks the bound fraction(*4*).

The goal of this work was to develop and apply a new class of dynamic reporters to allow for correlated light and electron (CLEM) imaging of protein interaction with molecular species not amenable to genetically encoded tags, such as DNA, RNA, or lipids. This strategy builds upon our click-EM method for labeling such macromolecular structures in cells with photosensitizing fluorophores(*5*). Metabolic substrates containing biorthogonal labels are incorporated into these macromolecules by endogenous enzymes(*6, 7*). A click reaction labels these azido or alkyne groups with a fluorescent dye for imaging and can also generate a localized osmiophilic precipitate visible to EM by photosensitization of DAB.

This new extension of this method, which we call “PS-PROMPT”, entails the chemical transfer of a photosensitizing fluorophore from a protein to a clickable metabolite that has been incorporated into cellular macromolecules or lipids by pulse chase. The requirement for close physical proximity of protein and macromolecule for photosensitizer transfer ensures only interacting fractions are so marked with untransferred removed before imaging. The PS-PROMPT probe has a modular structure composed of a protein-self labeling ligand e.g., a chloroalkane chain (HaloTag substrate, labeled HTL in Figure 1A), a linker of variable length (zigzag line, L in Figure 1A), a cleavable group such as disulfide bond (SS), a fluorescent photosensitizer dye (PS, Figure 1A, B) and a click-ready group (e.g., N3 or azido).

**Figure 1:**
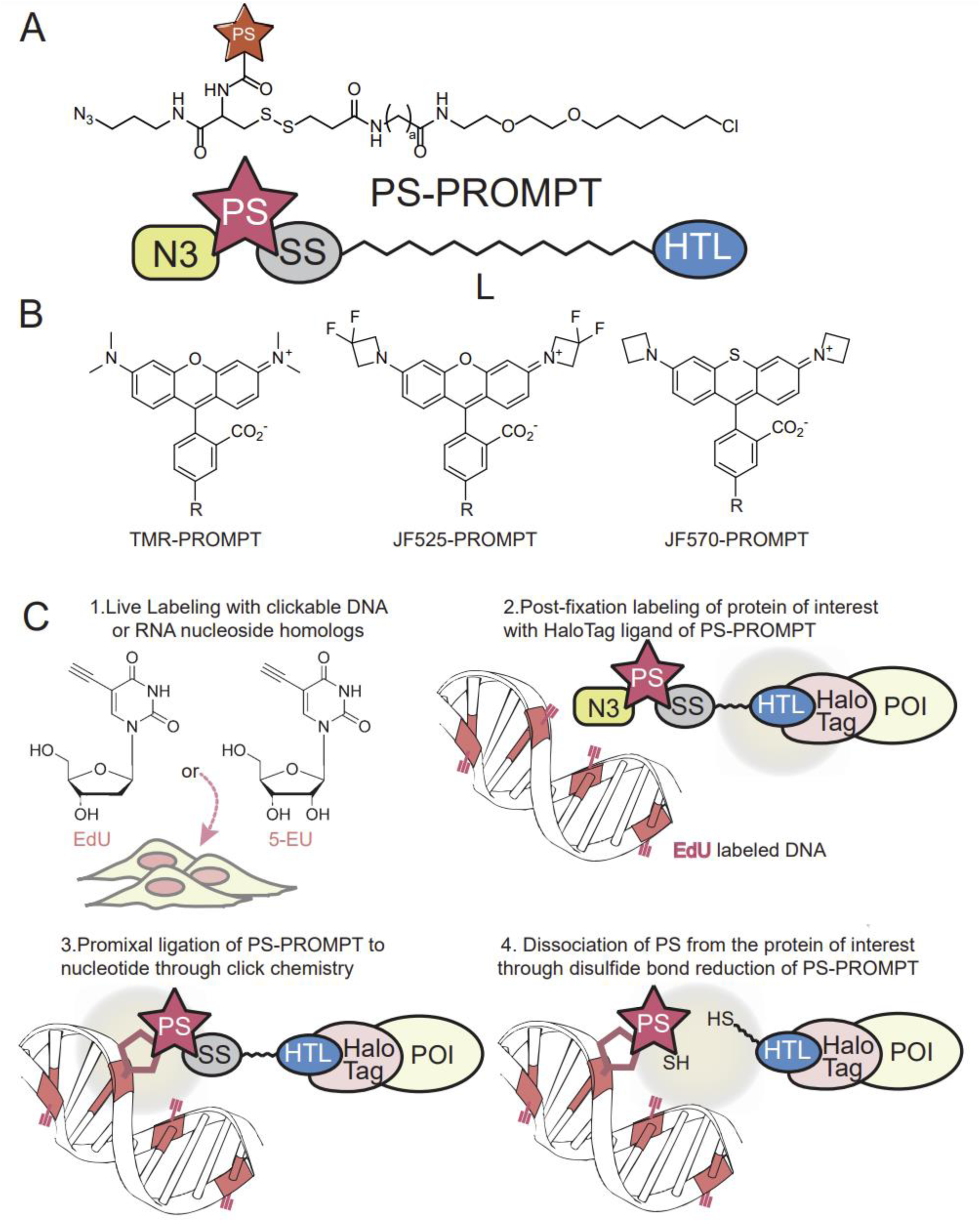
Principal components of a PS-PROMPT probe and the PROMPT procedure for detecting protein-DNA binding. A. Chemical structure and schematic of PS-PROMPT modules. N3: azido group; PS: Photosensitizers; SS: disulfide bond; HTL: HaloTag ligand, i.e., chloroalkane, a: methylene or polyethyleneglycol chain (L) of variable length. B. Chemical structures of the fluorescent photosensitizers in PROMPT probes. R: site of attachment in PROMPT probe. C. PROMPT Procedure overview for detection of binding between a protein of interest (POI) and DNA.

An example of using PS-PROMPT to detect interaction of a specific protein to DNA is depicted in Figure 1C. Cells expressing the DNA binding protein of interest fused to a HaloTag(*8*) protein are incubated with EdU (5-ethynyl-2’-deoxyuridine) to incorporate alkyne groups in newly synthesized DNA(*9*). Following fixation, the protein-HaloTag target is labeled with the PS-PROMPT probe. The probe is then covalently cross-linked by a click reaction(*9*) of the azido group to an alkyne group in bound DNA that is within the reach of its flexible linker. Unclicked probe is removed by reductive cleavage of the disulfide bond by dithiothreitol (DTT) and subsequent washes. Thus, the fluorescent photosensitizer is transferred from the protein to nearby DNA revealing the sites of interaction by LM through its fluorescence and by EM after photosensitization of DAB precipitation and subsequent osmification. In proof-of-principle tests, this new PS-PROMPT probe system revealed the interaction sites on DNA of the histone H2B fused with a HaloTag. Application of the method to the RNA binding protein, fibrillarin revealed its ultrastructural localization when bound to RNA.

## RESULTS

### Design and synthesis of PS-PROMPT

The PROMPT method requires the synthesis of a multimodal compound, generically named PS-PROMPT for PhotoSensitizer-PROMPT. The PS-PROMPT includes a chloroalkane group as a HaloTag ligand (HTL), an azido group (N3) compatible with click chemistry, a disulfide bond (SS) that can be cleaved by reduction, a linker of variable length (L), and a fluorescent photosensitizer (PS) (Fig. 1A). For application to correlative light and electron microscopy approaches, we chose the Janelia fluorophores, JF525 and JF570 as potent photosensitizers for DAB photooxidation(*10*), as well as tetramethylrhodamine (TMR) (*11*)(Fig. 1B). The fluorescence emission peaks of these dyes range from 549-599 nm (Fig. S1).

To test the functionality of each module of PS-PROMPT we first synthesized TMR-PROMPT (Fig. S2). 5(6)-carboxy-tetramethylrhodamine (TMR) was reacted with the 3-azidopropyl amide of S-trityl cysteine, prepared by reaction of Fmoc-NH-cys(S-Trityl)-OH and 3-azidopropyl-1-amine followed by removal of the N-terminal Fmoc. Cleavage of the trityl group and subsequent reaction of the thiol with the SPDP conjugate of HaloTag linker amine afforded TMR-PROMPT. JF570-PROMPT and JF525-PROMPT were synthesized (Fig. S3) by a similar procedure to TMR-PROMPT except the SPDP-HaloTag linker conjugate was reacted before final addition of the photosensitizer. Reaction of JF570-PROMPT and JF525-PROMPT with purified HaloTag protein in vitro was confirmed by LC-MS, giving the expected mass increase for the labeled protein.

### Validating the PS-PROMPT method

We first tested whether TMR-PROMPT can bind to HaloTag in tissue culture cells. For this, we expressed the histone H2B fused with HaloTag in U2OS cells and incubated them with TMR-HaloTag ligand (TMR-HTL) or TMR-PROMPT (Fig. 2A). TMR-HTL binds efficiently to HaloTag-H2B whether the incubation is on live cells or following aldehyde fixation, whereas TMR-PROMPT labels its target only when added after fixation. Because of this result and the potential for premature cleavage of the disulfide group by intracellular cysteines and glutathione, any PROMPT assay was performed following aldehyde fixation. The next set of experiments challenged the capability of dithiothreitol (DTT) to reduce the bound TMR-PROMPT disulfide bond and release the TMR fluorophore (Fig. 2B). Fixed U2OS cells transfected with HaloTag-H2B were incubated with TMR-HTL or TMR-PROMPT, then treated with 10 mM DTT and heavily washed. With or without DTT, cells incubated with TMR-HTL display a constant TMR fluorescent signal, verifying that DTT does not affect the fluorescent properties of TMR. However, cells treated with TMR-PROMPT and reduced with DTT, completely lose the nuclear TMR fluorescence, presumably by reductive cleavage of TMR-PROMPT and TMR release from HaloTag-H2B. Lastly, we verified the functionality of the azido group of TMR-PROMPT by performing Cu(I)-catalyzed azide-alkyne cyclization (CuAAC) click chemistry with TMR-PROMPT on EdU treated cells (Fig. 2C). Only when cells are labeled with EdU, is TMR fluorescence observed in the nucleus, and as expected, subsequent treatment with DTT on clicked TMR-PROMPT does not trigger TMR release.

**Figure 2:**
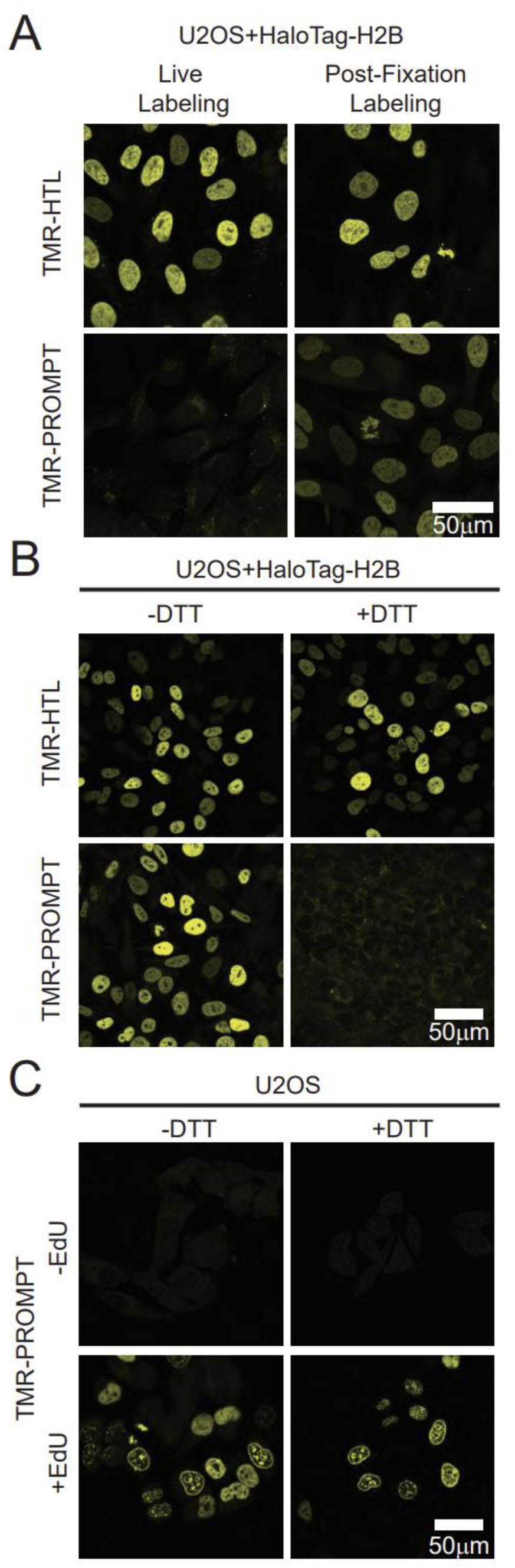
Testing functionality of the PROMPT probe modules. A. Confocal micrographs of U2OS cells transfected with HaloTag-H2B construct and incubated with 1 μM of TMR-HaloTag ligand (TMR-HTL) or 1 μM of TMR-PROMPT live for 1H or overnight at 4°C after 4% PFA fixation. B. Confocal micrographs of U2OS cells transfected with HaloTag-H2B construct and incubated overnight with 1 μM of TMR-HTL or 1 μM of TMR-PROMPT after 4% PFA fixation. Cells are subsequently treated with or without 10 mM DTT. C. Confocal micrographs of untransfected U2OS cells pretreated overnight with or without 5 μM of EdU. Following fixation in 4% PFA, the cells were labeled with TMR-PROMPT as above, covalently linked to incorporated EdU in DNA by click chemistry, and cells then treated with or without DTT as in B.

Next, we tested the feasibility of the PROMPT approach for CLEM using again as a model system the interaction between histone H2B with DNA. For CLEM, we used JF570-PROMPT, as JF570 is a potent photosensitizer(*10*). For the JF570-PROMPT procedure (Fig. 3A), following overnight incubation with 5 μM EdU, cells transfected with HaloTag-H2B are fixed with 4% paraformaldehyde (PFA) and 0.1% glutaraldehyde (GA) and briefly incubated with JF570-PROMPT in the presence of saponin to facilitate probe penetration. After washing, we verified that only HaloTag-H2B expressing cells display JF570 fluorescence in the nucleus, thereby demonstrating the covalent bond formation between JF570-PROMPT and HaloTag-H2B (Fig. 3B, top panel). Next, applying CuAAC click chemistry resulted in the reaction of the JF570-PROMPT immobilized on HaloTag-H2B with the nearest clickable substrate, an alkyne-uridinyl residue incorporated in DNA. After washing to remove the click chemistry solution and terminate the reaction, we treated the cells with DTT to cleave the disulfide link between JF570 and HaloTag-H2B. After intense washing to remove unclicked but cleaved JF570-PROMPT, we imaged the JF570 fluorescence (Fig. 3B, bottom panel). Gratifyingly, only in the condition where HaloTag-H2B is expressed in EdU-treated cells, do the majority of cells display a JF570 fluorescent signal above background in the nucleus (Fig. 3C). Interestingly, when comparing the JF570 fluorescence before the click chemistry and after the DTT treatment, we observed a significant reduction (5.2-fold) in mean intensity, suggesting that only a fraction of the JF570 bound to HaloTag-H2B is transferred to EdU (Fig. 3D).

**Figure 3:**
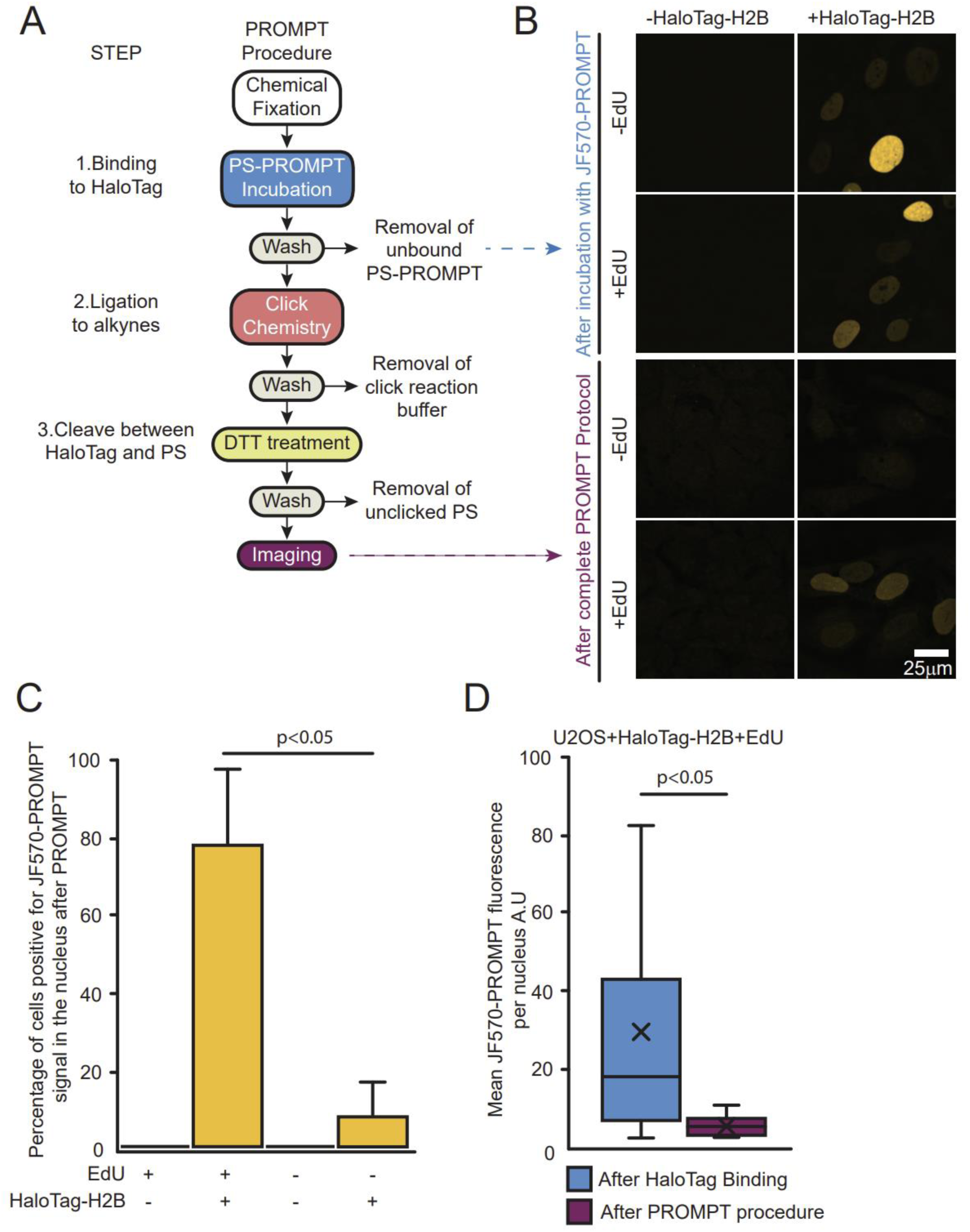
PROMPT protocol. A. Detailed schematic of the PROMPT protocol. B. Confocal micrographs of U2OS cells transfected with or without the HaloTag-H2B construct, pretreated overnight with or without 5 μM of EdU, and processed with the PROMPT protocol. In the top panel, the imaging is performed after JF570-PROMPT incubation and washing. In the bottom panel, the imaging is performed after the complete procedure, i.e., after DTT treatment and washing. C. Graph bars plot the average percentage of cells positive for JF570-PROMPT fluorescent signal after PROMPT protocol as in 2B bottom panel. D. Quantification of the mean JF570-PROMPT fluorescence signal per nucleus in cells expressing HaloTag-H2B and pretreated with EdU as in 2B. Imaging is performed after the JF570-PROMPT incubation or after the complete PROMPT procedure. Error bars are box-and-whiskers plots containing the mean (X), quartiles (box), and minimum and maximum observations (whiskers).

### PROMPT dependence on molecular distances in cellular ultrastructure

Considering the structural constraints on the PS transfer success rate, we reasoned that elongating the linker chain of PS-PROMPT (L in Fig. 1A) would increase its efficiency. We therefore added a PEG4 linker to TMR-PROMPT (Fig. 4A) and tested the TMR transfer efficiency. In cells expressing HaloTag-H2B and labeled with EdU, the TMR fluorescence intensity is significantly greater after PROMPT when using TMR-PEG4-PROMPT (Fig. 4B) compared to TMR-PROMPT. Elongating the probe not only compensates for unfavorable orientations of partners but also increases the radius in which the PS-PROMPT can find a clickable target.

**Figure 4:**
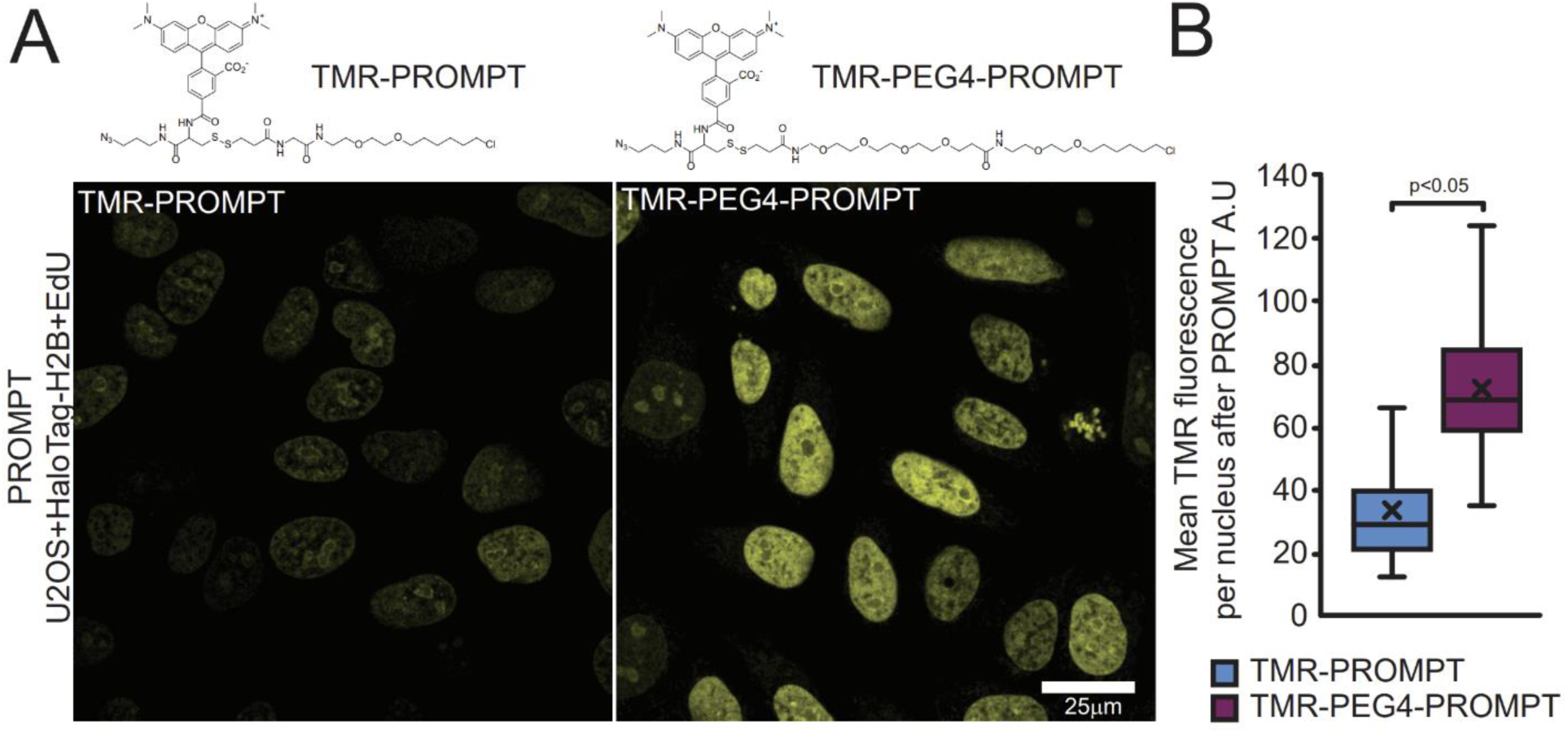
PROMPT for distance estimation. A. U2OS cells expressing HaloTag-H2B and pretreated with 5 μM EdU are processed with PROMPT using TMR-PROMPT probe or TMR-PEG4-PROMPT. A higher fluorescence signal is observed by confocal for TMR-PEG4-PROMPT compared to TMR-PROMPT. B. Quantification of the mean TMR fluorescence signal per nucleus in cells treated as 4A. Error bars are box-and-whiskers plots containing the mean (X), quartiles (box), and minimum and maximum observations (whiskers).

### Visualizing Histone H2B-DNA interaction in U2OS cells with PROMPT and CLEM

Next, we performed photooxidation of DAB through the illumination of the JF570-PROMPT after the PROMPT procedure in cells expressing HaloTag-H2B and labeled with EdU (Fig. 5A). After osmification, embedding, and sectioning, we verified the overall darkening of the correlated nucleus by electron microscopy. At higher magnification, we visualized a high concentration of puncta that were more opaque to electrons. To gain confidence regarding the identity of the puncta, we performed multi-tilt axis tomography, combining 1936 images of the same field of view taken at multiple angles(*12*). We obtained a volume of the isometric pixel size of 1.4 nm (Fig. 5B, Supplementary movie 1). The size and shape of some of the electron-dense puncta (highlighted by the arrowheads) are consistent with the expected nucleosomes.

**Figure 5:**
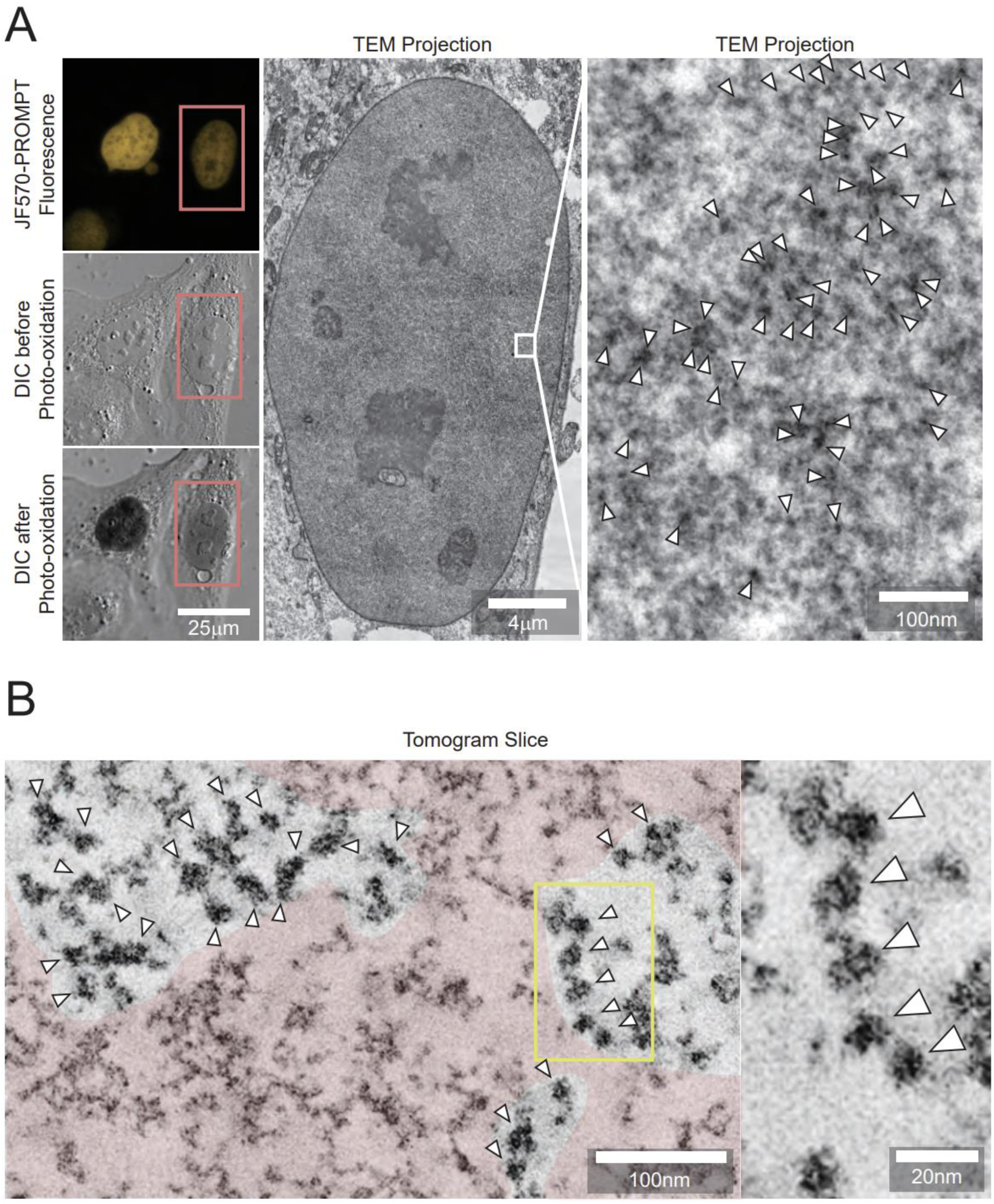
CLEM of histone H2B binding to DNA using PROMPT. A. The top left panel shows confocal, and DIC imaging (middle left) of EdU treated HaloTag-H2B expressing U2OS cells processed with JF570-PROMPT. The lower left image shows the amplitude of the DAB photooxidation by JF570-PROMPT. The center image shows an EM micrograph of the cell’s nucleus framed in the pink box in the left panels. The image on the right shows the enlargement of the area bounded in the white box in the center image. B. Both panels display an electron tomogram slice from a cell treated and photooxidized as in 5A. The image on the right is a close-up view of the area contained in the yellow box on the left panel. The pink shaded area highlights a region lacking the electron-dense puncta. For A. and B., arrowheads point at some of the electron-dense particles.

Interestingly, whereas some chromatin regions are enriched with these dark puncta, others are devoid of those (pink area, Fig. 5B). The lack of DAB precipitates indicates the failure of JF570-PROMPT transfer. The dilution of HaloTag-H2B by endogenous H2B, partial replacement of thymidines by EdU, and non-favorable orientations of the HaloTag active pocket and the clickable nucleotide bases within its reach could explain the inefficient transfer.

### Visualizing Fibrillarin-RNA interaction in U2Os cells with PS-PROMPT and CLEM

As we have demonstrated the feasibility of PROMPT to pinpoint protein-DNA binding partners by CLEM, replacing EdU with 5-Ethinyl Uridine (5EU) should make PROMPT amendable to the study of protein-RNA interaction. As a study system, we tested PROMPT on the interaction of fibrillarin with RNAs. As a component of the C/D box small nucleolar ribonucleoproteins, fibrillarin is involved in the 2’-O-methylation of rRNAs(*13, 14*). We transfected U2OS cells with HaloTag-Fibrillarin and labeled overnight newly synthesized RNAs with 5EU. After PROMPT using JF570-PROMPT, the JF570 fluorescent signal is only detected in cells treated with both HaloTag-Fibrillarin and 5EU (Fig. 6A, B). Similar observations were made using JF525-PROMPT (Fig. S4) where JF570 has been replaced with the yellow photosensitizer, JF525. Completing the protocol with DAB photooxidation by JF570, the extent of DAB polymerization was dependent on the JF570 fluorescence intensity as expected (Fig. 6C). After correlation, we imaged at high magnification the nucleolus and surroundings by EM (Fig. 6D, E). As expected, we observed a high density of dark particles in the nucleolus (cyan dotted line boundaries) and at the Cajal bodies (purple dotted line boundaries). Interestingly, we also detected evidence of fibrillarin transferring PROMPT to RNAs at discrete puncta of a few nanometers throughout the nucleoplasm (Fig. 6E, arrowheads) and in 200 to 250 nm granules (Fig 6D, white arrows) of unknown identity. Electron tomography not only confirms the preponderance of fluorophore transfer sites in the nucleolus and the Cajal body but also highlights Fibrillarin and RNA interaction clusters at the surface of these organelles (Fig. 7, Supplementary movie 2). Furthermore, the 3D distribution of the fibrillarin-RNA complexes throughout the nucleus is made visible. In summary, the PROMPT method confirmed the known interaction of fibrillarin with rRNAs for their methylation and revealed potential functions in the nucleoplasm.

**Figure 6:**
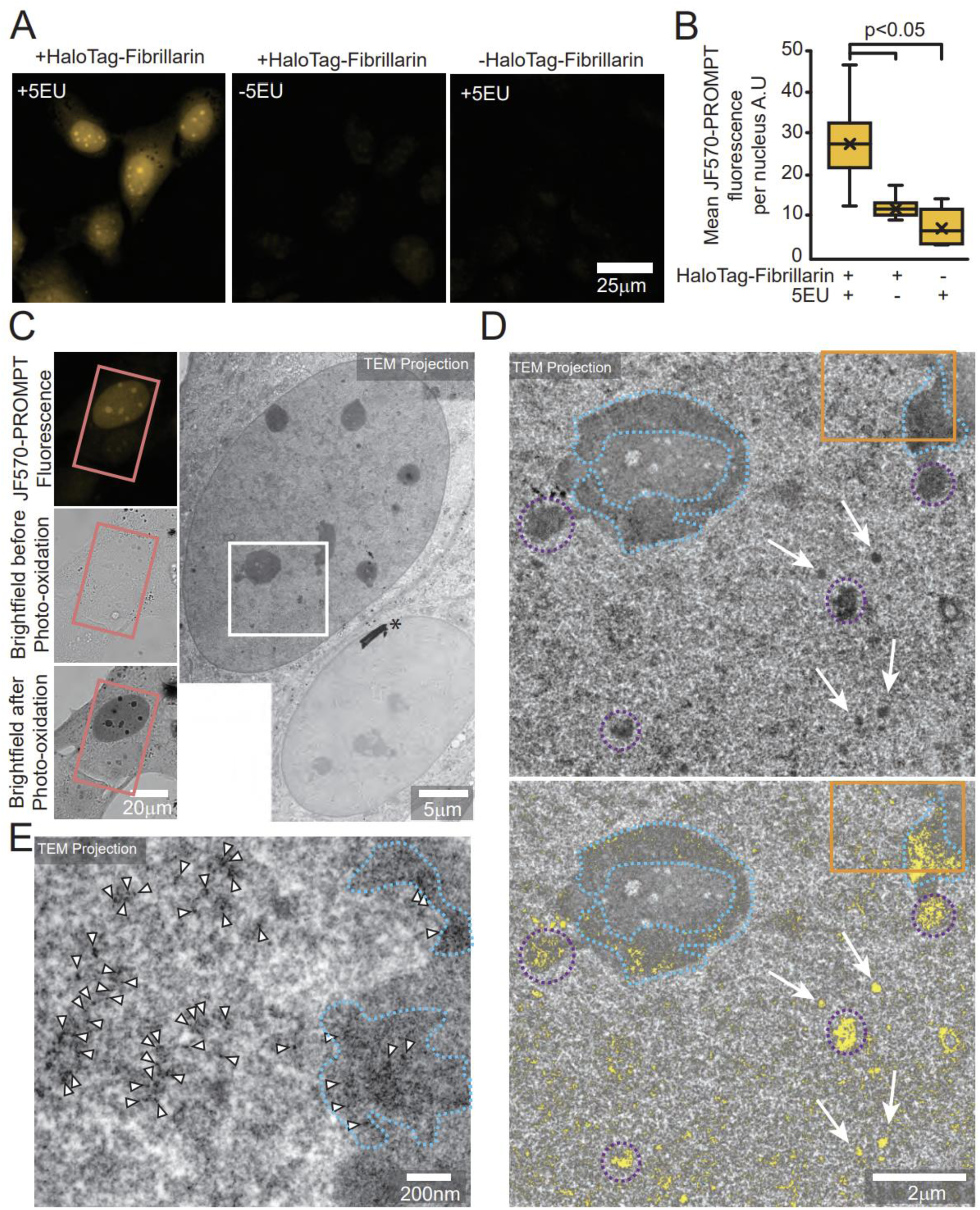
CLEM of Fibrillarin binding to RNA using PROMPT. A. Confocal micrographs of U2OS cells transfected with or without the HaloTag-Fibrillarin construct, pretreated overnight with or without 1 mM 5EU, and processed with the PROMPT procedure. B. Quantification of the mean JF570-PROMPT fluorescence signal per nucleus in cells treated as in 6A. Error bars are box-and-whiskers plots containing the mean (X), quartiles (box), and minimum and maximum observations (whiskers). C. DAB photooxidation by JF570-PROMPT after the PROMPT procedure in U2OS cell transfected with HaloTag-Fibrillarin construct and treated with 1 mM 5EU. The left micrographs show the fluorescence and the brightfield images pre and post photooxidation. The image on the right shows the electron microscopy micrograph of the cells shown on the left and framed in the pink box. The asterisk points at a speck of dust on the resin section. D. Electron microscopy micrograph of the area of interest in the white box in C. The top image shows the original acquisition, and the bottom one is a composite image including the original TEM micrograph layered with the DAB polymers extracted by thresholding and colorized in yellow. The purple dotted lines circle the Cajal bodies. The white arrows point at 250 nm diameter granules of unknown identities. E. Electron microscopy micrograph of the area of interest in the orange box in D. Arrowheads point at some of the electron-dense DAB polymers. D, E. The cyan dotted line surrounds darker areas of the nucleolus.

**Figure 7:**
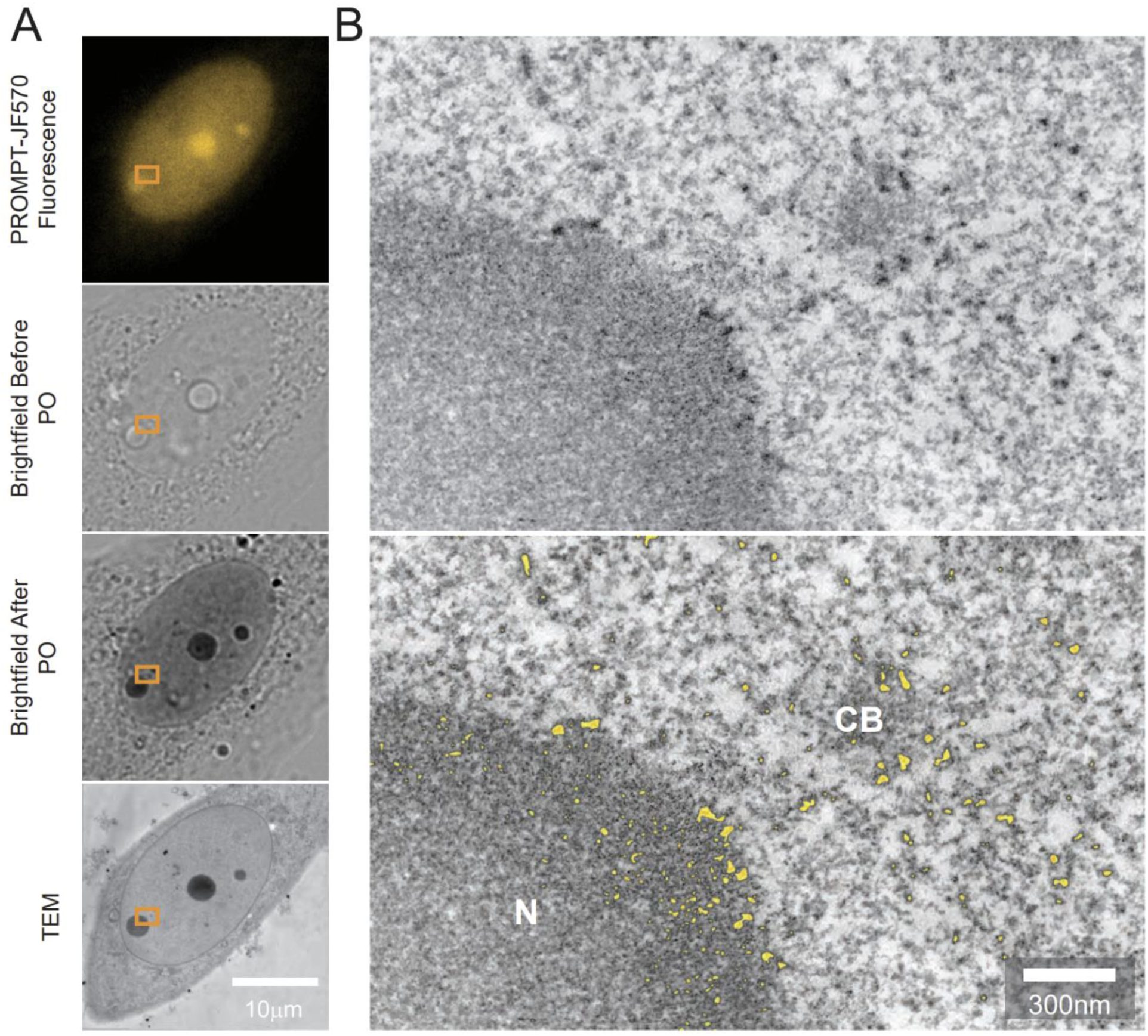
Electron Tomography of Fibrillarin/RNA PROMPT. A. Confocal micrographs of U2OS cells transfected with the HaloTag-Fibrillarin construct, pretreated overnight with 1 mM 5EU, and processed with the PROMPT procedure. Brightfield images are acquired before and after pho-oxidation (PO) of DAB by JF570-PROMPT. The lowest image shows the corresponding photo-oxidized cell by transmission electron microscopy. The orange box delineates the area acquired by electron tomography in 7B. B. Both panels display a minimum projection of 8 consecutive electron tomogram slices from the cell treated and photo-oxidized in 7A. In the lowest image, the most electron dense areas have been segmented by thresholding and colorized in orange. N: Nucleolus; CB: Cajal Body.

## DISCUSSION

In this study, we developed PROMPT, a new class of molecular probes and method for the visualization of the interaction of proteins with DNA or RNA demonstrating the detection by correlated light and electron microscopy of the binding of DNA with the histone H2B and RNAs with fibrillarin. The chains of nucleosomes visualized using PROMPT for histone H2B and DNA interaction validates the specificity and high-resolution images obtainable with the method. In addition to verifying the known nuclear localization of fibrillarin, PROMPT reveals its interactions with RNAs, presumably rRNAs and small nucleolar RNAs(*15*). In addition, the method reveals a previously undescribed nuclear but non-nucleolar fraction of fibrillarin is also associated with RNAs suggesting fibrillarin remains associated with rRNAs upon ribosome maturation, on its entry to or exit from the nucleolus.

The PROMPT procedure itself is relatively simple and, except for the probe, does not require specialized chemicals or equipment. It can potentially be applied to visualize the interaction of any HaloTag fusion protein with cellular components that can incorporate clickable metabolites such as proteins, nucleotides, sugars, lipids, enzymatic inhibitors, etc. The method is amenable for spatial-temporal studies using pulse-chase to detect lifetimes of HaloTag fusion proteins interaction with newly synthesized macromolecules. Conversely, to measure binding of new HaloTag fusion proteins by pulse chase would require blocking old protein with a conventional HaloTag ligand, followed by its omission for a defined time before fixation. Because the PS-PROMPT probe is not incorporated in live cells, only newly synthesized proteins would be accessible to post-fixation labeling by the current PS-PROMPT probe.

The inability of PS-PROMPT to label live cells probably reflects a low membrane permeability that could be improved by replacing the polar amide bonds with more hydrophobic groups. Additionally, to prevent reduction of the disulfide bond of the PS-PROMPT probe in the reducing environment of the cytoplasm, replacement with a photocleavable linker(*16*) or periodate-cleavable vicinal diol-containing linker(*17*) would be required. One potential indication of this phenomenon is the residual fluorescence in the oxidizing environment of the endoplasmic reticulum on live labeling with TMR-PROMPT (Fig. 2A) that may reflect reaction of its cleaved disulfide group(*18*) with endogenous ER thiols, and therefore the non-specific anchoring of the fluorophore. The PS-PROMPT cleavage system was inspired by the cleavable SNAP ligands used to study the dynamics of endocytosis(*19*). In this context, the impermeable probe is always on the luminal side of the cell and therefore not sensitive to cytoplasmic reduction.

The partial lack of transfer of the fluorophores between H2B and DNA, while potentially revealing some biological phenomena, may also highlight certain limitations of the method. These limitations are the coexistence of HaloTag-H2B and endogenous H2B, the partial replacement of thymidines by EdU(*20*), and steric and distance constraints in transfer reaction. Incomplete HaloTag binding and click reaction may also be at play here. Alternative methods for the click chemistry might improve the transfer efficiency(*21*).

The choice of the protein fused to HaloTag could facilitate the identification and subcellular localization of specific DNA or mRNA sequences in cells. In combination with dCas9-HaloTag with a specific guide RNA(*4*), one could target only the binding fraction of dCas9 to the chosen DNA sequences. While relatively weak, the signal should be detectable by removal of the excess, unbound fraction of labeled dCas9-HaloTag that loses fluorescence by the PROMPT method. Similar approaches can be applied to mRNA using MS2 protein fused with HaloTag(*22*). The PROMPT method should reduce this background signal and highlight only the mRNA fused with the MS2 sequences and bound to the MS2 protein.

The PROMPT probe is composed of different functional modules, and the modification of one or several modules broadens the range of potential applications of the method. Replacement of the HaloTag ligand module by SNAP or CLIP ligands (benzyl guanine and benzyl cytosine respectively)(*23*) expands the spectrum of protein targeting systems and enables simultaneous PROMPT for multiple proteins. Replacing alkyne groups incorporated with metabolic intermediates such as EdU with SNAP or CLIP ligands containing alkynes, would potentially allow CLEM applications of the PROMPT method for protein-protein interaction. Probably the most exciting applications of PROMPT may come from the replacement of the module that is being transferred. For instance, the photosensitizer can be replaced by a wide range of fluorophores, including the ones compatible with super-resolution, such as JF549 or JF646(*24*). An interesting modification would be to add an additional fluorophore (of non-overlapping spectral properties or part of a FRET donor-acceptor pair) to the HaloTag side of the disulfide bond, so it is retained when PROMPT transfer does not occur. In this way, one could potentially determine both the total protein pool with this fluorophore and the bound fraction with the transferred fluorophore. By incorporating a biotin group, PROMPT could also enable pulldown of binding partners for sequencing and/or mass spectrometry, in addition to imaging their interaction. Finally, PROMPT can potentially serve as a molecular ruler and highlight structural changes of the protein *in situ* that positively or negatively affect the transfer rate.

## MATERIAL and METHODS

### Cell culture, 5EU and EdU labeling, transfection

U2OS cells were cultured on 35 mm MatTek dishes (MatTek Corp) in DMEM supplemented with 10% FBS at 5% CO2. On reaching 70% confluency and two days before the experiment, cells are transfected with 0.5 μg of DNA with lipofectamine 3000 (ThermoFisher Scientific) following the manufacturer protocol. If needed, cells are incubated overnight in culture medium with 5 μM of 5-ethynyl 2’-deoxyuridine (EdU, #1149, Click Chemistry Tools) diluted from a 10 mM stock in DMSO. For 5-EU (5-ethynyl uridine) incorporation, 200 mM 5-EU (#1261, Click Chemistry Tools) in DMSO is diluted to 1 mM in culture medium and incubated with cells overnight.

### DNA constructs

For the construct of HaloTag-H2B (JH1348), the H2B sequence was amplified from pminiSOG-H2B-6 (*25*) and substituted EGFP in pHaloTag-EGFP (addgene #86629)(*26*) through the In-Fusion cloning method. For the construct of HaloTag-Fibrillarin (JH1239), the Fibrillarin sequence was amplified from pEGFP-C1-Fibrillarin (addgene #26673)(*27*) and substituted EGFP in pHaloTag-EGFP (addgene #86629) through the In-Fusion cloning method.

### Synthesis of TMR-PROMPT and TMR-PEG4-PROMPT **(Fig. S2)**

#### NH2-cys(S-Trt)-CO-NH-(CH2)3-N3 (2)

Fmoc-NH-cys(S-Trt)-OH, (**1**) (19.8 mg, 33.8 μmol) and HATU (14.1 mg, 37.2 μmol) were dissolved in dry DMF (100 µL) in a plastic screw cap tube and 3-azidopropyl-1-amine (3.7 μL, 37.2 μmol, Click Chemistry Tools) followed by DIEA (13 μL, 74.4 μmol) were added with mixing. The reaction mix turned yellow and LC-MS revealed complete reaction in 30 mins; ES-MS (m/z) [M+ Na]^+^ for C40H37N5NaO3S, 690.25; found 689.3. Piperidine (20 μL, 0.2 mmol) was added and the solution evaporated under high vacuum after 1 h, dissolved in DMSO, separated by RP-HPLC and lyophilized to give **2** as a white solid. Yield, 14.3 mg, 76%. ES-MS (m/z) [M]^+^, [M+ Na]^+^ for C25H27N5OS, 446.2, 468.2; found 446.1, 468.1.

#### 5(6)-TMR-CONH-cys(SH)-CO-NH-(CH2)3-N3 (4)

5(6)-Carboxytetramethylrhodamine, 5(6)-TMR-CO2H (1.35 mg, 3.14 μmol, Novabiochem) and TSTU (1.3 mg, 4.4 μmol) were dissolved in dry DMSO (25 μL) with TEA (0.96 μL, 6.9 μmol) and kept at room temperature. Reaction was complete in 30 min (by LC-MS) and then added to a solution of NH2-cys(S-Trt)-CO-NH-(CH2)3-N3 (**2**) (2.0 mg, 3.6 μmol) in DMSO (10 μL) with NMM (1 μL, 9.1 μmol) and kept at room temperature overnight when LC-MS revealed complete reaction. After acidification with HOAc (2 μL), the desired product, **3** was isolated by RP-HPLC and lyophilized to a red solid. Yield, 2.0 mg (74%) ES-MS (m/z) [M]^+^ for C50H48N7O5S, 858.3; found 858.3. The trityl group was removed by dissolving the product (1.89 mg, 2.2 μmol) in TFA:H2O:Triisopropylsilane:Ethanedithiol (92.5:2.5:2.5:2.5 v/v, 0.5 mL) for 30 mins, evaporation under high vacuum, purification by RP-HPLC and lyophilization to give **4** as a red solid. Yield, 0.9 mg (66%) ES-MS (m/z) [M]^+^ for C31H34N7O5S, 616.2; found 616.1.

#### SPDP-HaloTag linker

HaloTag linker amine (*28*) (3.0 mg, 13.5 μmol) and SPDP (4.7 mg, 15 μmol) were dissolved in dry DMSO (50 μL) and NMM (3.3 μL, 30 μmol) added. LC-MS revealed reaction was complete after overnight when the reaction mixture was neutralized with HOAc (5 μL) and the product was purified by RP-HLPC (and lyophilized to a colorless oil. Yield, 3 mg (54%) ES-MS (m/z) [M]^+^ for C18H30ClN2O3S2, 421.1; found 421.1.

#### TMR-PROMPT; 5(6)-TMR-CONH-cys(S-S-(CH2)2CONH-HaloTag ligand)-CO-NH-(CH2)3-N3 (5)

A solution of 5(6)-TMR-CONH-cys(SH)-CO-NH-(CH2)3-N3 (**4**) in DMSO (100 μL, 6.25 mM measured by absorbance in 0.1 M HCl in 95% ethanol using εmax 95000 M^-1^cm^-1^ at 554 nm, 0.626 μmol) was mixed with SPDP-HaloTag linker (20 μL, 49 mM in dry DMSO, 0.98 μmol) and NMM (1 μl, 10 μmol) added. After 1 h, HOAc (5 μL) was added and the desired product, (**5**) purified by RP-HPLC to give a colorless oil. Yield, 0.42 mg (72%) ES-MS (m/z) [M]^+^ for C44H58ClN8O8S2, 925.4; found 925.4.

#### SPDP-PEG4-HaloTag linker

Solutions of HaloTag linker amine (*28*) (3.0 mg, 4.5 μmol,) in dry DMSO (90 μL) and SPDP-PEG4-NHS (2.5 mg, 4.5 μmol, Quanta Biodesign) in dry DMSO (90 μL) were mixed and NMM (1 μL, 10 μmol) added. LC-MS revealed reaction was complete after 4h and the product was used without further purification. ES-MS (m/z) [M]^+^ for C29H51ClN3O8S2, 668.3; found 668.3.

#### TMR-PEG4-PROMPT; 5(6)-TMR-CONH-cys(S-S-(CH2)2CONH-PEG4-HaloTag ligand)-CO-NH-(CH2)3-N3 (6)

5(6)-TMR-CONH-cys(SH)-CO-NH-(CH2)3-N3 (**4**) (3 mL of 0.25 mM in DMSO; measured as above) was added to the solution of SPDP-PEG4-HaloTag Linker (15 μL of 50 mM, 0.75 μmol) and NMM (15 μl, 150 μmol) added. After 2h, LC-MS indicated reaction was complete, HOAc (50 μL) was added, the product, (**6**) purified by RP-HPLC and lyophilized. Yield, 0.7 μmol (by absorbance in 0.1M HCl in 95% ethanol using εmax 95000 M^-1^cm^-1^ at 554 nm) after dissolving in dry DMSO (100 μL). ES-MS (m/z) [M]^+^ for C55H79ClN9O13S2, 1172.5; found 1172.4.

### Synthesis of JF525-PROMPT and JF590-PROMPT **(Fig: S3)**

#### Fmoc-NH-cys(SH)-CO-NH-(CH2)3-N3 (7)

Fmoc-NH-cys(S-Trt)-OH (**1**) (59 mg, 100 μmol) and HATU (42 mg, 110 μmol) were dissolved in dry DMF (200 µL) in a plastic screw cap tube and 3-azidopropyl-1-amine (11 μL, 110 μmol, Click Chemistry Tools) followed by DIEA (38 μL, 220 μmol) were added with mixing. The reaction mix turned yellow and LC-MS revealed complete reaction in 30 mins; ES-MS (m/z) [M+ Na]^+^ for C40H37N5NaO3S, 690.3; found 690.2. The reaction mixture was evaporated to yellow oil and trityl group removed by dissolving in TFA-H2O-Triisopropylsilane-Ethanedithiol (92.5/2.5/2.5/2.5 v/v, 1 mL) and kept at room temperature for 2h. Following evaporation to an oily solid, the desired product was purified by RP-HPLC, and lyophilized to give (**7**) a white solid. ES-MS (m/z) [M]^+^ for C21H24N5O3S, 426.2; found 426.1.

#### NH2-cys(S-S-2-(CH2)2CONH-HaloTag ligand)-CO-NH-(CH2)3-N3

Fmoc-NH-cys(SH)-CO-NH-(CH2)3-N3 (**7**) (1.25 μmol, 50 μL of 25 mM solution in dry DMSO) was mixed with SPDP-HaloTag linker (1.25 μmol, 25 μL of 50 mM solution in dry DMSO) and NMM (2.5 μL, 25 μmol) added. LC-MS revealed complete reaction after 20 min, ES-MS (m/z) [M]^+^ for C34H48ClN6O6S2, 735.3; found 735.3. Piperidine (20 μL) was added and the solution evaporated after 5 mins. The residue was dissolved in DMSO and HOAc (5 μL), product **8** isolated by RP-HPLC, lyophilized and dissolved in dry DMSO (75 μL). ES-MS (m/z) [M]^+^ for C19H38ClN6O4S2, 513.2; found 513.2.

#### JF570-PROMPT; Janelia Fluor 570-CONH-cys(S-S-(CH2)2CONH-HaloTag ligand)-CO-NH-(CH2)3-N3 (9)

The product**, 8** from the previous step was reacted with JF570-NHS ester (0.4 μmol, 8 μl of 50 mM solution in dry DMSO) and NMM (10 μmol, 1 μl) for 2 days at room temp, the product **9** isolated by RP-HPLC, lyophilized and dissolved in DMSO (100 μL) to give a 1.4 mM solution, measured by εmax 100000 M^-1^cm^-1^ in 0.1 M HCl in 95% ethanol at 574 nm. ES-MS (m/z) [M]^+^ for C46H58ClN8O7S3, 965.3; found 965.3.

#### JF525-PROMPT; Janelia Fluor 525-CONH-cys(S-S-(CH2)2CONH-HaloTag ligand)-CO-NH-(CH2)3-N3 (10)

Prepared as for JF570-PROMPT. Measured by εmax 80000 M^-1^cm^-1^ in 0.1 M HCl in 95% ethanol at 532 nm. ES-MS (m/z) [M]^+^ for C46H54ClF4N8O S ^+^,1021.3; found 1021.3.

#### Reaction with HaloTag protein

JF525-PROMPT or JF570-PROMPT (0.5 μL of 1 mM solution in DMSO) was added to HaloTag protein (2.5 μL of 100 μM solution in PBS) diluted in 100 mM Na·MOPS pH 7.2, and kept at room temperature for 2h. Acetic acid (1 μL) was added and analyzed by LC-MS using a PLRP-S 1000A column (8 μm, 50 x 2.1 mm, Agilent) eluting with linear 20-60% ACN-water with constant TFA (0.05%) in 16 min.

JF570-PROMPT: ES-MS (m/z) [M]^+^ for adduct with HaloTag protein (loss of HCl), C46H57N8O7S3, (965.3-35.98) 929.3; for deconvoluted masses of JF570-PROMPT:HTP complex, 35729.0 minus HTP, 34798.6; found 930.4.

JF525-PROMPT: ES-MS (m/z) [M]^+^ for adduct with HaloTag protein (loss of HCl), C46H53F4N8O8S2, 985.3; for deconvoluted masses of JF525-PROMPT:HTP, 35783.5 minus HTP, 34798.6, found 984.9.

### PROMPT Method

After EdU or 5EU labeling, transfected cells are washed with culture medium and fixed for 2 minutes with room temperature fixative (4% EM grade paraformaldehyde (Electron Microscopy Sciences #19202) + 0.1% EM grade glutaraldehyde (Ted Pella, #18426) in 0.1 M Hepes (Sigma-Aldrich #H3375) pH7.4 + 2 mM CaCl2) followed by a 1-hour incubation with 4°C fixative. Next, the cells are washed three times with PBS and incubated for 10 minutes with PBS+ 10 mM glycine. The cells are then incubated for 30 minutes with 200 nM of PROMPT probe in PBS+0.1% BSA (Sigma-Aldrich #A8022) +0.1% saponin (Sigma-Aldrich #S4521), then washed five times with PBS+0.1%BSA+0.1%saponin, and five times with PBS. For the click chemistry step, the cells are incubated for 2 times 30 minutes with 50 mM Hepes pH7.4 + 100 mM NaCl + 2 mM CuSO4 + 10 mM of freshly prepared sodium ascorbate (Sigma-Aldrich #A7631). The reaction is terminated with ten washes of PBS. For the disulfide bond reduction, cells are incubated three times 10 minutes with freshly prepared 10 mM DTT (Sigma-Aldrich #D9779) in PBS. Released photosensitizers are removed with five washes of PBS+0.1%BSA+0.1%saponin and five washes with PBS.

### Imaging and Photooxidation

If not for EM, cells were imaged in PBS with an inverted confocal microscope Leica SPEII. For photooxidation, cells were imaged in 2.3 mM DAB (Sigma-Aldrich #D8001) in 0.1 M cacodylate (Sigma-Aldrich #C0250) buffer pH 7.4 with the same microscope. DAB was photo-oxidized by JF570-PROMPT under illumination with a 63X objective for 12 minutes using the 575/25 line (310mW) of a Spectra 7 LED light source (Lumencor). Then samples were washed with 0.1 M cacodylate buffer pH 7.4 and processed for EM.

### Image analysis

Fluorescence intensities were measured using ImageJ. Mean intensity were calculated from more than 100 cells from 3 independent experiments.

### Resin embedding, sectioning, and TEM imaging

After photo-oxidation, samples were post-fixed for 30 minutes with 1% osmium (Electron Microscopy Sciences #19150) + 1.5% potassium ferrocyanide in 0.1 M cacodylate buffer pH 7.4 at 4°C. Then the samples were dehydrated in 1-minute successive 4°C baths of increasing ethanol concentration (20%, 50%, 70%, 90%, 3 x 100%), and incubated for 1-hour in 50%ethanol/50% durcupan resin (Sigma-Aldrich #44611, #44612, #44613, #44614). Samples were then incubated 3 x 2-hours in 100% durcupan mixture, and the resin was polymerized for 48H at 60°C. At the photo-oxidized areas, blocks of hardened resin were cut off from the dish and mounted onto dummy blocks using glue. The glass was popped up by water infiltration, and the blocks were trimmed around the cells of interest to shape a block face of approximatively 400 μm wide and 200 μm tall. Block faces were imaged with an upright light microscope at 10X to facilitate correlation. Ribbons of 70nm thick sections were produced from these blocks and collected on glow-discharged formvar coated slot grids (Electron Microscopy Sciences #FF2021-Cu-ET). Areas of interest were imaged at 120 KeV with an FEI Tecnai Spirit transmission electron microscope.

### Multi tilt axis electron tomography

For electron tomography, 250 nm thick sections were produced and collected onto Luxel grids (Luxel, #LuxFilmTEM C-S-M-L), then incubated for 20 seconds with 20% 10 nm gold fiducials (Ted Pella#15703-20) + 0.05% BSA in water, blotted, and dry. The grids with sections were then imaged with an upright light microscope at 10X. The grids were loaded onto a rotation holder (Fischione Instruments, model 2040). After correlation, a four tilt-axis series (one tilt-series: −60° to +60°, one image every 0.25°) at different orientation (0°, 45°, 90°, 135°) of the region of interest was acquired using an FEI Titan Halo operated at 300 KeV and a DE64 camera, while controlled by SerialEM(*29*). The entire set of 1936 acquired images were aligned and reconstructed using TxBR package(*30, 31*).

## ACKNOWLEDGMENTS

This project was supported by the following grants, R01 GM086197 (SRA/DB), NSF2014862 (MHE), R01AG081037 (MHE), and R01GM138780 (MHE).

We thank John Ngo for helpful discussions, Hiroyuki Hakozaki for technical assistance, and Luke D. Lavis for graciously providing us with JF525 and JF570 dyes.

---

**Figure S1:**
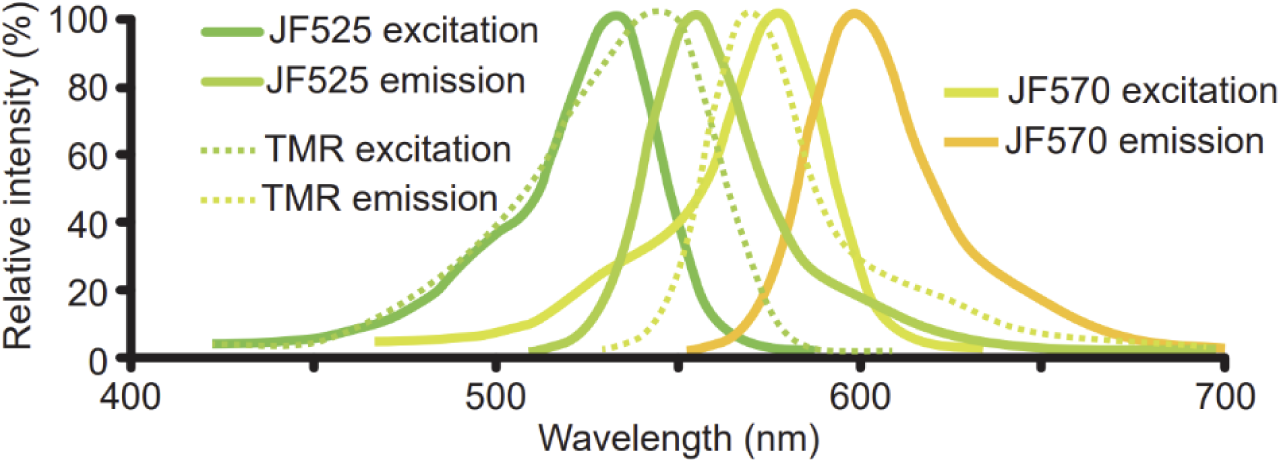
Photosensitizer spectra used in PROMPT.

**Figure S2:**
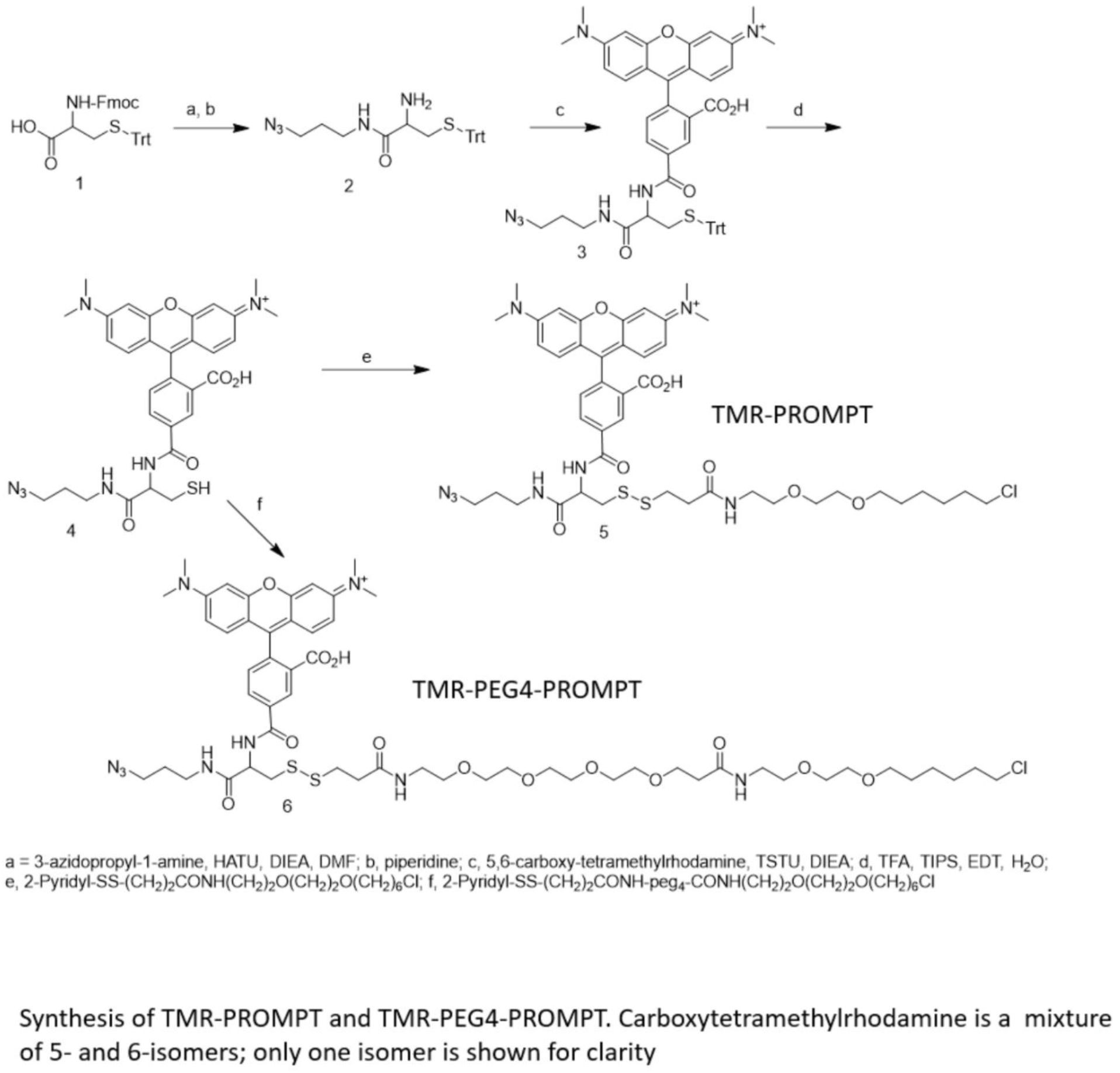
Synthesis of TMR-PROMPT and TMR-PEG4-PROMPT

**Figure S3:**
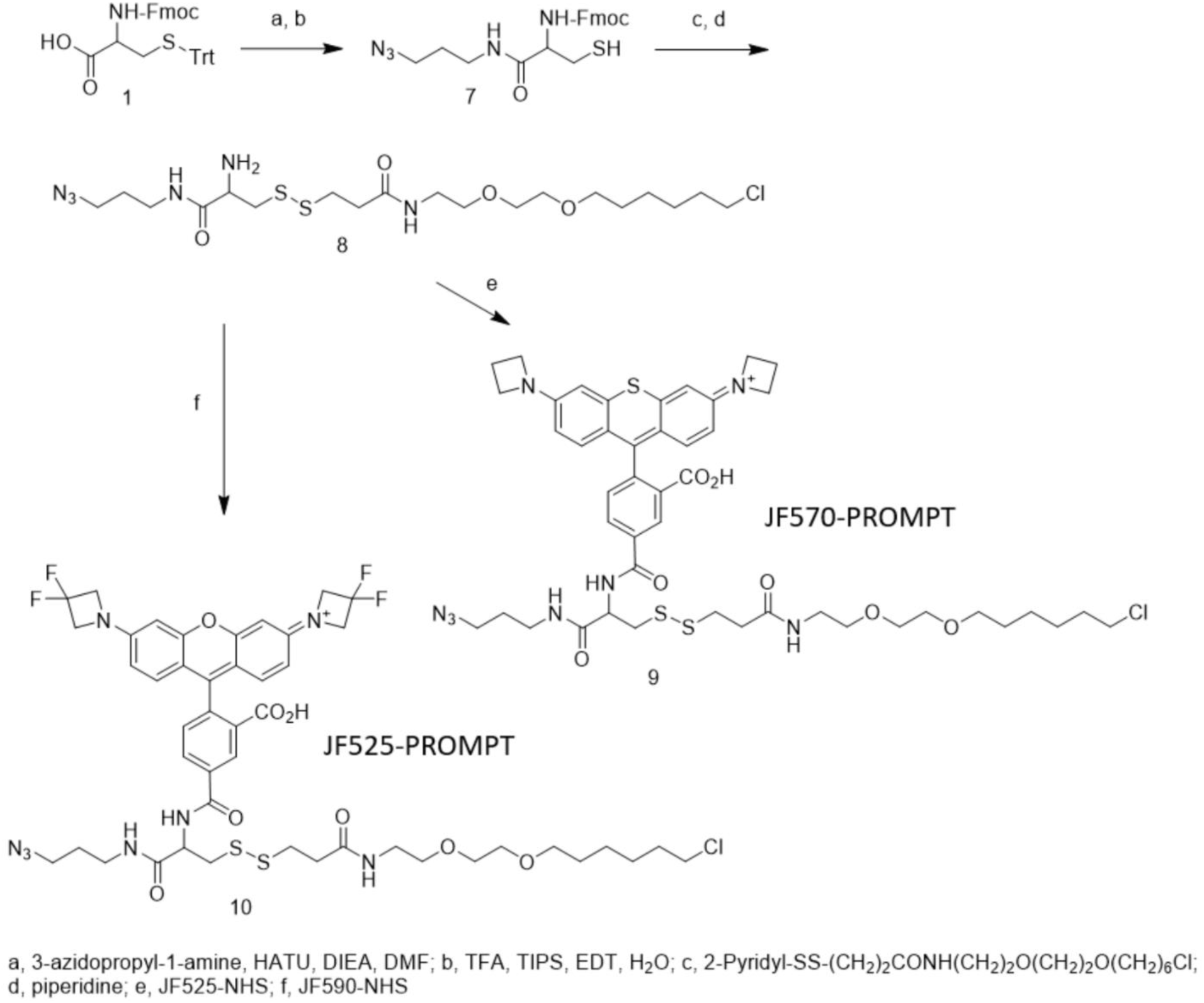
Synthesis of JF570-PROMPT and JF525-PROMPT

**Figure S4:**
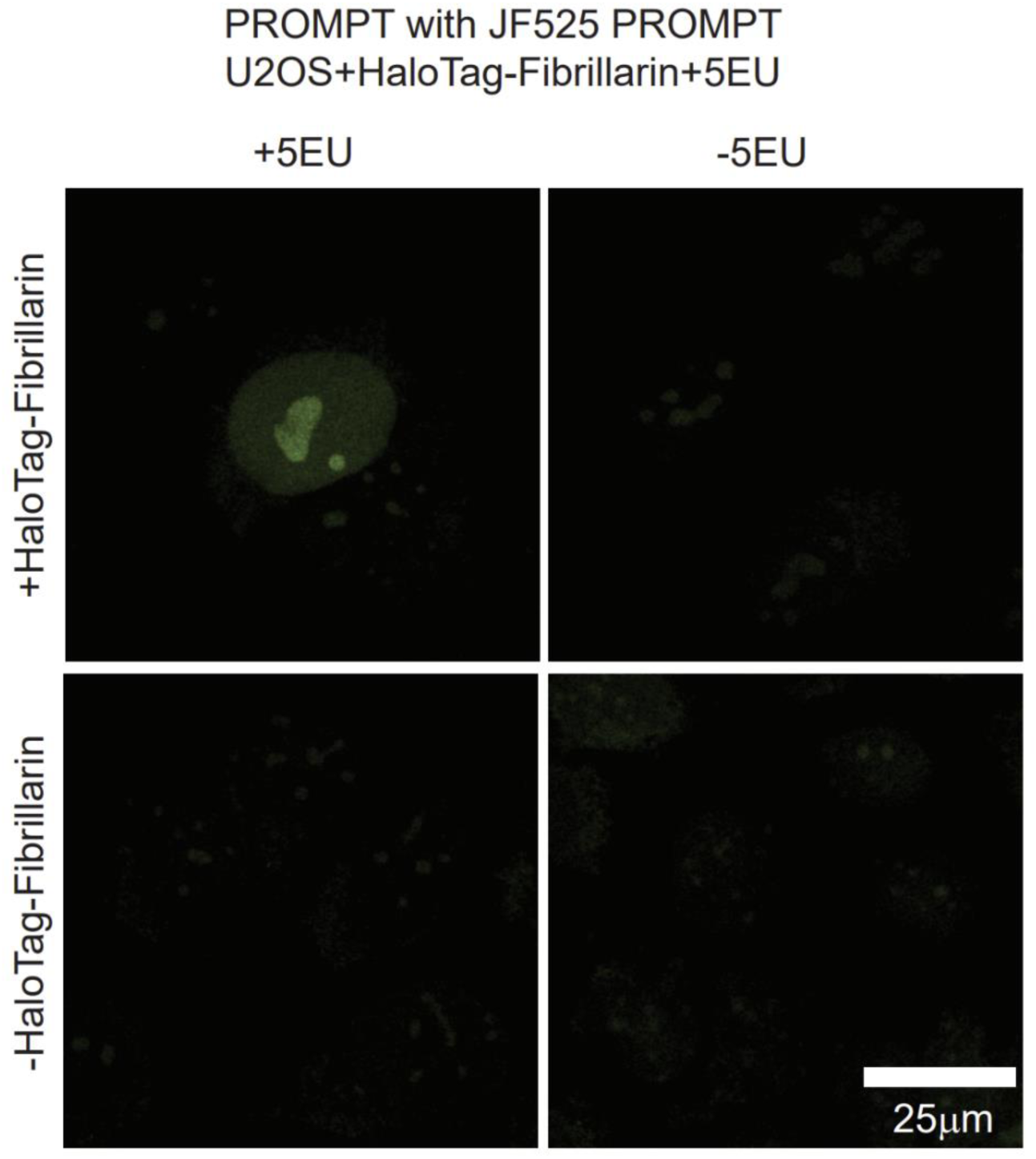
Fibrillarin/RNA PROMPT with JF525-PROMPT. Confocal micrographs of U2OS cells transfected with or without the HaloTag-Fibrillarin construct, pretreated overnight with or without 1mM 5EU, and processed with the PROMPT procedure using JF525-PROMPT.

**Supplementary movie 1**: Multi-tilt axis electron tomogram of a U2OS cell expressing HaloTag-H2B pretreated overnight with 5uM of EdU, processed with the PROMPT procedure and DAB photo-oxidized. The total tomogram thickness is 250 nm.

**Supplementary movie 2**: Multi-tilt axis electron tomogram of a U2OS cell expressing HaloTag-Fibrillarin pretreated overnight with 1uM of 5EU, processed with the PROMPT procedure and DAB photo-oxidized. The total tomogram thickness is 250 nm.

## REFERENCES

1. B. T. Bajar, E. S. Wang, S. Zhang, M. Z. Lin, J. Chu, A Guide to Fluorescent Protein FRET Pairs. Sensors. 2016 (10.3390/s16091488).

2. D. W. Piston, G.-J. Kremers, Fluorescent protein FRET: the good, the bad and the ugly. Trends in Biochemical Sciences 32, 407–414 (2007).

3. W. Deng, X. Shi, R. Tjian, T. Lionnet, R. H. Singer, CASFISH: CRISPR/Cas9-mediated in situ labeling of genomic loci in fixed cells. Proceedings of the National Academy of Sciences 112, 11870–11875 (2015).

4. B. Chen et al., Dynamic Imaging of Genomic Loci in Living Human Cells by an Optimized CRISPR/Cas System. Cell 155, (2013).

5. J. T. Ngo et al., Click-EM for imaging metabolically tagged nonprotein biomolecules. Nature Chemical Biology 12, 1–10 (2016).

6. C. Y. Jao, A. Salic, “Exploring RNA transcription and turnover in vivo by using click chemistry,” (2008).

7. A. Salic, T. J. Mitchison, A chemical method for fast and sensitive detection of DNA synthesis in vivo. Proceedings of the National Academy of Sciences 105, 2415–2420 (2008).

8. G. V. Los et al., HaloTag: A Novel Protein Labeling Technology for Cell Imaging and Protein Analysis. ACS Chemical Biology 3, 373–382 (2008).

9. V. V. Rostovtsev, L. G. Green, V. V. Fokin, K. B. Sharpless, A Stepwise Huisgen Cycloaddition Process: Copper(I)-Catalyzed Regioselective “Ligation” of Azides and Terminal Alkynes. Angewandte Chemie International Edition 41, 2596–2599 (2002).

10. T. C. Binns et al., Rational Design of Bioavailable Photosensitizers for Manipulation and Imaging of Biological Systems. Cell Chemical Biology 27, 1063–1072.e1067 (2020).

11. V. Liss, B. Barlag, M. Nietschke, M. Hensel, Self-labelling enzymes as universal tags for fluorescence microscopy, super-resolution microscopy and electron microscopy. Nature Publishing Group 5, 17740–17740 (2015).

12. H. D. Ou et al., ChromEMT: Visualizing 3D chromatin structure and compaction in interphase and mitotic cells. Science 357, eaag0025 (2017).

13. J. P. Aris, G. Blobel, “cDNA cloning and sequencing of human fibrillarin, a conserved nucleolar protein recognized by autoimmune antisera (evolutionary conservation/sderoderma/RNA-binding motif),” Proc. Nati. Acad. Sci. USA (1991).

14. U. Rodriguez-Corona, M. Sobol, L. C. Rodriguez-Zapata, P. Hozak, E. Castano, Fibrillarin from Archaea to human. Biology of the Cell 107, 159–174 (2015).

15. Z. Kiss-László, Y. Henry, J.-P. Bachellerie, M. Caizergues-Ferrer, T. Kiss, Site-Specific Ribose Methylation of Preribosomal RNA: A Novel Function for Small Nucleolar RNAs. Cell 85, (1996).

16. S. V. Wegner, O. I. Sentürk, J. P. Spatz, Photocleavable linker for the patterning of bioactive molecules. Scientific Reports 5, (2016).

17. R. J. Smith, R. A. Capaldi, D. Muchmore, F. Dahlquist, Crosslinking of ubiquinone cytochrome c reductase (complex III) with periodate-cleavable bifunctional reagents. Biochemistry 17, 3719–3723 (1978).

18. I. Braakman, H. Hoover-Litty, K. R. Wagner, A. Helenius, Folding of influenza hemagglutinin in the endoplasmic reticulum. Journal of Cell Biology 114, (1991).

19. N. Jaensch, I. R. Corrêa, R. Watanabe, Stable Cell Surface Expression of GPI-Anchored Proteins, but not Intracellular Transport, Depends on their Fatty Acid Structure. Traffic 15, (2014).

20. T. E. Creighton, Conformational restrictions on the pathway of folding and unfolding of the pancreatic trypsin inhibitor. Journal of Molecular Biology 113, (1977).

21. A. R. Frand, C. A. Kaiser, Ero1p Oxidizes Protein Disulfide Isomerase in a Pathway for Disulfide Bond Formation in the Endoplasmic Reticulum. Molecular Cell 4, (1999).

22. E. Bertrand et al., Localization of ASH1 mRNA Particles in Living Yeast. Molecular Cell 2, 437–445 (1998).

23. A. Gautier et al., An Engineered Protein Tag for Multiprotein Labeling in Living Cells. Chemistry & Biology 15, (2008).

24. J. B. Grimm et al., A general method to improve fluorophores for live-cell and single-molecule microscopy. Nature Methods 12, (2015).

25. X. Shu et al., A genetically encoded tag for correlated light and electron microscopy of intact cells, tissues, and organisms. PLoS Biology 9, (2011).

26. M. Ebner, I. Lučić, T. A. Leonard, I. Yudushkin, PI(3,4,5)P3 Engagement Restricts Akt Activity to Cellular Membranes. Molecular Cell 65, 416–431.e416 (2017).

27. D. Chen, S. Huang, “Nucleolar Components Involved in Ribosome Biogenesis Cycle between the Nucleolus and Nucleoplasm in Interphase Cells,” The Journal of Cell Biology (2001).

28. R. C. Strauch et al., Reporter protein-targeted probes for magnetic resonance imaging. J Am Chem Soc 133, 16346–16349 (2011).

29. D. N. Mastronarde, in Journal of Structural Biology. (Academic Press Inc., 1997), vol. 120, pp. 343–352.

30. S. Phan et al., TxBR montage reconstruction for large field electron tomography. Journal of Structural Biology 180, 154–164 (2012).

31. S. Phan et al., 3D reconstruction of biological structures: automated procedures for alignment and reconstruction of multiple tilt series in electron tomography. Advanced Structural and Chemical Imaging 2, (2016).

